# MicroNucML: A machine learning approach for micronuclei segmentation and the refinement of nuclei-micronuclei relationships

**DOI:** 10.1101/2025.09.20.677550

**Authors:** Yukai Wang, Nadejda B. Boev, Ulises O. Garcia, Kate M. MacDonald, Shane M. Harding, Sushant Kumar

## Abstract

Micronuclei (MN) are structures containing small fragments of DNA, arising from mitotic errors or failed DNA repair attempts. Therefore, MN serve as markers of genomic instability and are typically quantified either manually or through threshold-based methods, which can be tedious and inaccurate, leading to varying degrees of success and throughput. By employing a two-phase labeling approach that utilizes polygon and brush segmentation, along with refinement using SAM2, we developed a high-quality MN segmentation tool. Subsequent data augmentation, which captured heterogeneity in image quality and color diversity, enabled us to train a generalizable Mask-RCNN model optimized for small object detection, achieving state-of-the-art performance in MN detection. Finally, we applied our model to immunofluorescence data obtained from cell lines exposed to DNA damage conditions to gain biological insights into MN dynamics and their role in inducing genome instability. In summary, this work establishes an accessible resource for systematically studying genome instability with significantly greater fidelity and sensitivity, enabling insights into damage biology that were previously unresolved.

## Background

Micronuclei (MN) are small, extranuclear structures that encompass nuclear contents, including DNA and DNA-binding proteins such as histones, which are collectively encapsulated by an envelope^1^. MN can arise from mitotic errors or inadequately resolved DNA breaks, thus serving as markers for chromosomal instability (CIN)^2,3^. For instance, CIN can be exemplified by chromothripsis (i.e., a shattered chromosome), which is frequently observed in cancer cells^4,5^. One major cause of chromothripsis is the chromosomal sequestration in MN. Downstream to MN formation, MN can rupture and activate the innate immune system via the cGAS-STING pathway, subject to the detection of its contents in the cytoplasm^6,7^. In practice, MN are characterized to assess cancer risk^8^, implicated when screening for aneuploidy in embryos^9^, and when evaluating the genotoxicity of environmental compounds^10,11^. Consequently, there has been a longstanding interest in identifying, quantifying, and characterizing MN within various biological and clinical contexts.

Common experimental approaches for MN visualization through microscopy include directly staining DNA or fluorescently labeling histones. Subsequently, manual counting or threshold-based segmentation can be performed using software such as ImageJ to enumerate MN^12^. While manual annotation is accurate^13^, it is time-consuming and lacks scalability for high-throughput experiments. Moreover, due to the small size of MN, the segmentation quality may vary across labelers. In contrast, thresholding, which utilizes watershed-based algorithms, can be scaled up; however, it requires manual input of parameters for object size and intensity, making it highly dependent on image quality and/or experimental conditions^14^. Importantly, watershed strategies are impartial to the objects detected. For instance, apoptotic fragmentation or nuclear blebbing may be incorrectly included due to size-based object detection. Therefore, a machine learning (ML) strategy may address the scalability and variation challenges in MN detection and segmentation.

To date, several ML models have emerged for the segmentation of nuclei, such as DeepCell^15^, CellPose^16^, and StarDist^17^. However, these models are primarily trained on images captured under normal conditions with undamaged DNA. Consequently, they currently lack the ability to identify structures like MN, which are relevant for CIN scoring. Previous work for MN detection includes the development of a pre-trained ML model, NucRec, which classifies a nucleus as either having or being devoid of MN^18^. Although useful, this model was not trained to count or segment MN in large and busy images. In contrast, the model, MicroNuclAI, can count MN and nuclear buds (NBUDs) in complex images to produce a CIN score^19^. However, NBUDs are structurally distinct from MN; for example, NBUDs remain tethered to the main nucleus, can originate from distinct events, and may have differing fates^20,21^. Furthermore, MicroNuclAI only provides counts, rather than segmentations. Therefore, this limits the features that can be extracted from these structures, which are helpful for further characterization of MN. As such, there is a need to develop a tool that can scale, accurately segment MN, and be robust against experimental and technical variation in complex images.

Given the utility of MN identification, we aimed to develop an open-source model and tool that generates high-quality MN masks. We implemented a collaborative framework to efficiently label MN with experts, resulting in over 4000 manually segmented MN. We trained and optimized a Mask Region-based Convolutional Neural Network (Mask-RCNN) model, MicroNucML. We evaluated the model with particular attention to MN size and its generalizability across variations in image color and quality. Overall, our tool demonstrated state-of-the-art performance with high precision and recall for MN counting and segmentation. Furthermore, our model is color-invariant; therefore, it can be applied across a range of visualization techniques, including images in various colors, such as gray, blue, green, or red. These strategies can be used to visualize MN with nuclear dyes, such as DAPI, or fluorescent proteins, including GFP and mCherry. We then applied MicroNucML to live-cell images captured in a series of biological studies to characterize MN formation following DNA damage. We included an experiment using cGAS-tagged constructs to examine the dynamics of MN rupture, along with a large-scale study to measure MN production over time. Additionally, our tool features a safety mechanism to filter out objects suspected of apoptosis and, most importantly, integrates nuclei and MN segmentation. This distance-based strategy can thus discriminate between nuclei that produce MN and those that do not. The array of features captured by our tool provides deeper biological insights for CIN, addressing a pressing need for both clinicians and biologists investigating the fundamental processes driving genome instability.

## Results

### Description of the workflow for MN and nuclei segmentation

Our method utilizes a supervised machine learning framework that integrates manually labeled MN and nuclei segmentation **(Figure 1A)**. The framework begins with microscopy images, which visualize DNA via H2B-GFP-based fluorescence assays. The novelty of our method lies in precise MN segmentation, conducted through a two-phase approach: polygon segmentations using entire images, and brush segmentations using a filtered subset of image tiles **(Figure S1)**. We rationalized that this strategy could improve label efficiency by reducing the search space and easing the segmentation burden. As such, we performed manual labeling across large images (1400×1000 pixels) with polygon segmentations for both nuclei and MN, using LabelStudio^22^. These large images were then used to produce non-overlapping tiles (224×224 pixels), which either contained or excluded MN. These annotated images were then employed to train a classification model using ResNet101.

**Figure 1.**
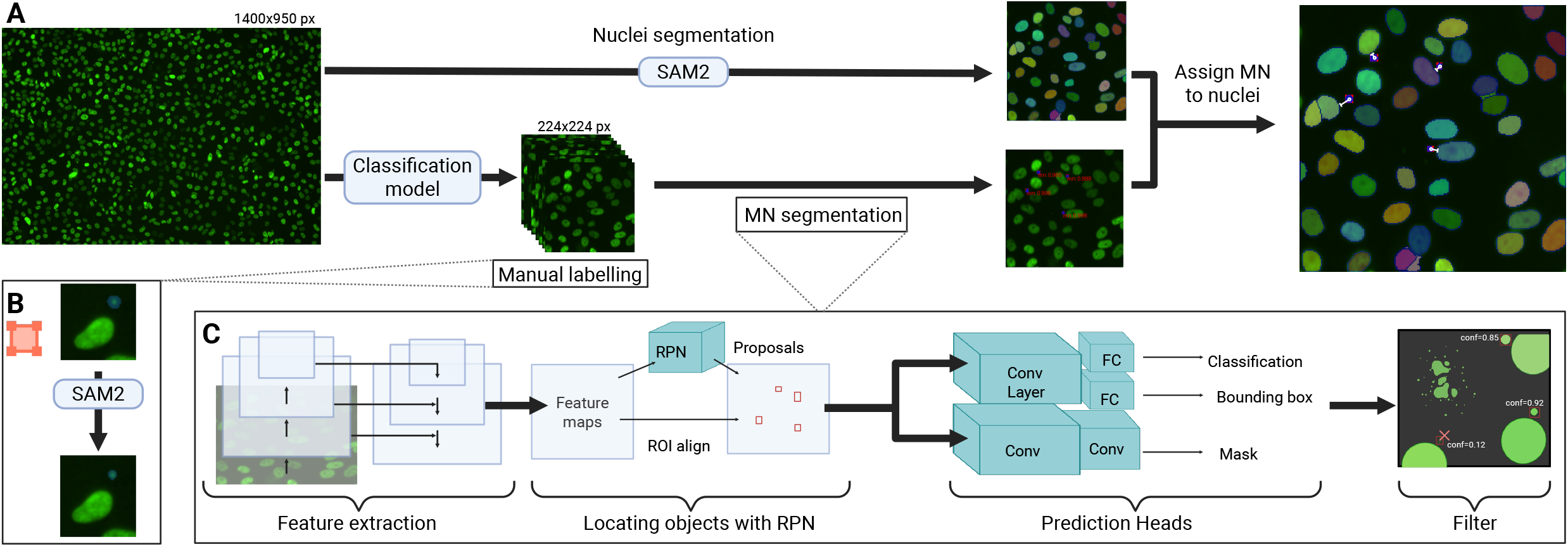
Overview of MicroNucML model framework for micronuclei (MN) and nuclei segmentation. **(A)**. Schematic of the input to output. H2B-GFP images from live-cell microscopy are the input. Nuclei segmentation is canonically performed with SAM2, whereby nuclei identified as similar in size to detected MN are filtered out. Alternatively, users can provide previously generated nuclei masks. In tandem, tiles with MN were classified using a binary classifier, then manually labelled with a brush tool. MN segmentations were used to train a Mask RCNN model. The final stage involves assigning MN to their parent nuclei using a distance-based approach. For each MN, the centroid is identified, whereby the most proximal nuclear edge is assigned as the parent nucleus. The end-user receives an output file listing the index of the parent nuclei, MN locations, MN sizes, and the model’s confidence for each segmented MN, along with nuclei features. **(B)** Brush segmentations produced with LabelStudio were further refined. The labellers’ segmentations were used as a point prompt by SAM2 to improve ground truth quality. **(C)** The Mask RCNN model begins with feature extraction via the optimized backbone, ResNet-50. A five-level feature map is then fed into a region proposal network (RPN). The model is then further optimized by varying the intersection over union (IOU) requirements and the number of anchors. The model is trained to minimize the classification, box regression, and mask logit losses. The final stage includes filtering out MN with poor confidence and removing proposed MN suspects of apoptosis.

This binary classification model was subsequently deployed to identify tiles with MN across full-sized images, which were then manually segmented using a brush tool. During the labeling process, we observed over-drawing with the brush tool; therefore, we utilized the Segment Anything Model (SAM2)^23^, a foundation model, to refine the segmentation masks **(Figure 1B)**. Notably, SAM2 is effective without fine-tuning and can be scaled. These masks were then employed to train a series of ML models for benchmarking MN segmentation. To address variability in image quality and color, we implemented a range of data augmentation strategies. Based on our systematic benchmarking, we found Mask-RCNN (**Figure 1C**) to be the optimal model for MN segmentation. Mask-RCNN models include a stage for region proposal, followed by object classification. During model optimization, we experimented with several backbones, which vary in their input data and architecture, alongside a feature pyramid network (FPN) using ResNet-50, Swin Transformer, and ResNeXt101. We refined the region proposal network (RPN) to enhance the detection of small and variable objects, such as MN. After identifying sufficiently confident MN segmentations, we evaluated the models. Most importantly, we investigated the impact of MN size on evaluation, using intersection-over-union (IOU) and scale-adaptive IOU (SIOU)^24^. Briefly, IOU compares the overlap between the ground truth and predicted bounding boxes. However, IOU is size invariant, meaning smaller objects are more harshly penalized for small errors in size or position. Therefore, we used IOU and SIOU when assessing the model using precision, recall, mean average precision (mAP), F1 score, and inference time.

Next, we observed that cells displaying an apoptotic-like morphology can present MN-like structures, indicating DNA fragmentation or blebbing. Although labelers avoided segmenting such objects, we expect users may provide images outside the distribution of the training images. Therefore, as a safety measure to protect against inaccurate classification, we included a post-processing feature that allows users to filter out predictions in regions with excessively high MN density. Following MN segmentation, we used SAM2 with customized point prompts to segment nuclei. We discarded segmentations of nuclei that appeared similar in size to the population of MN in the image. Finally, we aimed to assign MN to “parent” nuclei, thereby demarcating nuclei presumed responsible for MN production. We employed a distance-based approach, defining the parent as the most proximal nuclei based on the distance between the MN’s centroid and the nuclei’ edge.

### Performance evaluation of the trained Mask-RCNN for MN detection and segmentation

The availability of high-quality labeled data is essential for training a supervised ML model. However, manually segmenting entire images, particularly with closed polygons, is a labor-intensive task for label generation. Therefore, we adopted a two-step strategy for training our model. First, we reduced the visual search space for labeling by extracting smaller regions of interest (ROI) from the original images. Second, we applied a simple method (brush application in LabelStudio) to facilitate segmentation of images with ROI (i.e., presence of MN) with a single click. We trained a binary classification model using polygon-labeled images with a ResNet-101 architecture to predict whether a given image consists of ROIs. With five-fold cross-validation (CV), we achieved an average accuracy of 0.889. The final fine-tuned model demonstrated an accuracy, precision, and recall of 0.916, 0.805, and 0.991, respectively **(Figure S2)**.

We applied the trained binary classifier to filter images with putative MN. We then manually segmented 2,413 tiles using LabelStudio’s brush tool. We used SAM2’s refined segmentations as the ground truth. We found that MN are small, with a mean refined segmentation size of 27.2**±**19.4 pixels (at 0.625 µm/pixel) among the training dataset **(Figure S3)**. Therefore, we rationalized that our model architecture and final evaluation metrics should be suitable for objects of this size. We used the tiles to train a series of segmentation models, holding out 20% of the labeled tiles for testing. We began by comparing the single-stage and two-stage detectors, You Only Look Once (YOLO) and Mask-RCNN. When evaluating with an IOU threshold of 0.5, we found that the Mask-RCNN models significantly outperformed the YOLO models **(Figure 2A**). Next, we focused on optimizing the Mask-RCNNs by testing various RPN strategies and backbone architectures. Initially, by utilizing models with a ResNet-50 backbone, we optimized the number of anchors and IOU requirements within the RPN during training. As expected, we discovered that the model with anchor sizes customized for smaller objects (4, 8, 16, 32, 64), along with a lower IOU (0.5), performed exceptionally well. With these optimized parameters, we then focused on the model backbones using ResNet-50, Swin Transformer, and ReXt-101. We found that ResNet-50 achieved the best performance, with detection precision, recall, and mAP@50 values of 0.89, 0.91, and 0.96, respectively, at a confidence level of 0.7 and IOU of 0.5. We also observed satisfactory inference speed performance, with a processing time of 3.3 seconds using an NVIDIA T4 GPU **(Figure S4)** and 0.79 seconds with an A100 GPU. The optimized ResNet-50 model serves as the core module of our MN detection tool (microNucML).

**Figure 2.**
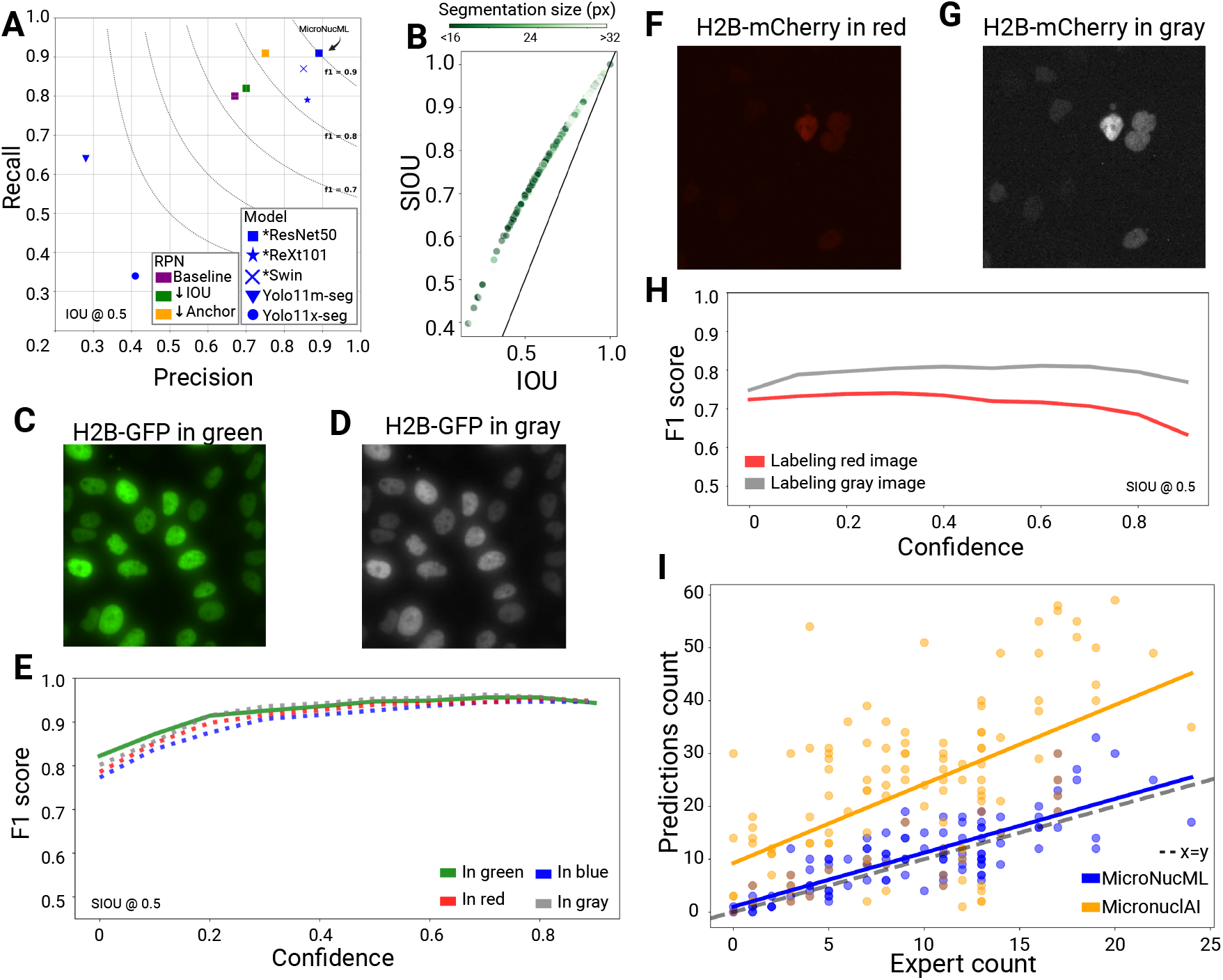
Performance evaluation and generalizability of the MicroNucML model. **(A)** Precision-Recall curve depicting the evaluation of MN segmentation on the held-out H2B-GFP testing dataset, at an IOU of 0.5. Trained models include those with a Mask-RCNN and YOLO architecture. Markers in blue demonstrate the final optimized models, as per their architecture and backbone. The superior model, a Mask RCNN with a ResNet-50 backbone, is highlighted with a black arrow. Markers in purple, green, and orange were derived during ablation studies. The dotted curves depict F1-scores. **(B)** A scatterplot comparing the IOU and scale-adaptive IOU (SIOU), as per testing the MN refined segmentation size. The colour gradient demonstrates MN size. **(C-E)** Evaluation of the model’s invariance to colour was conducted using green test images **(C)**, along with manipulated red, blue, and gray images **(D). (E)** As such, upon fixing the SIOU threshold to 0.5, MN confidence and F1 scores were captured across the four colour conditions. **(F-H)** An independent H2B-mCherry dataset was similarly manipulated, in red **(F)** to gray **(G). (H)** In this case, these poorer quality images were segmented separately into red and gray. A line graph depicts F1 scores in response to variations in MN confidence. **(I)** A scatterplot comparing expert and predicted MN counts from 60 H2B-GFP unseen images. Predictions and linear regressions were produced using MicroNucML MN counts (in blue) and MicronuclAI chromosomal instability scores (in orange). A black dotted diagonal line is shown at y=x.

Next, we examined the relevance of MN size in relation to model evaluation. When considering standard IOU, poor performance arises among reasonably sized objects due to errors in prediction size and/or location. However, for small objects, errors of even a few pixels can lead to poor IOU. Therefore, we investigated whether the model’s errors stem from imprecision in segmentation or if the evaluation metric is ill-suited for our task. Upon comparing the refined and predicted segmentation size of MN in the training dataset, we found very strong pixel-wise concordance (*β*=0.961, ρ=0.943, p-value=1.13e-296; **Figure S5A**). Next, to measure transformations between labels and predictions, we computed their Euclidean distance and found a non-significant relationship between the refined MN size and the shifts (*β*=-0.001, ρ=-0.054, p-value=0.175; **Figure S5B**). Therefore, we concluded that predictions are of high quality, both when predicting MN size and position; however, evaluation appears to be unfairly affected by MN size. Thus, we opted to use SIOU as our alternative evaluation metric. This strategy rescales IOU specifically for small objects and aligns well with human judgments of performance^24^. We observed an increase in performance when using SIOU instead of IOU, which was unique among the smallest MN **(Figures 2B and S5C-D)**.

Notably, our initial model was trained solely on H2B-GFP images, which may impact its generalizability to other fluorescence-tagged or stained nuclei. Therefore, we applied data augmentation strategies to increase the diversity of colors and image quality during training. Subsequently, we systematically evaluated the model’s generalizability. First, to test color invariance, we used the held-out H2B-GFP images **(Figure 2C)**, which had image qualities equivalent to those used for training. We adjusted the images’ color to gray **(Figure 2D)**, red, and blue to replicate the visual conditions used with mCherry constructs or DAPI/Hoechst staining. We found that even when segmentation confidence varies, performance remains unchanged based on color **(Figures 2E & Figure S6)**.

Next, we assessed the influence of variations in color and image quality on model performance. We used an independent set of H2B-mCherry images (n=308), which exhibited notably lower quality upon inspection **(Figure S7A)**. For instance, we observed increased background fluorescence and/or nuclei appearing out of focus. Importantly, the refined segmentation sizes of MN did not differ between H2B-mCherry and H2B-GFP images (Mann-Whitney U test (MW), p=0.614; **Figure S7B**). However, upon quantifying image sharpness, we found that the H2B-mCherry images were significantly less sharp compared to the H2B-GFP images (p=4.637e-116; **Figure S7C**). This issue posed challenges during manual segmentation, and we noted minimal improvements from SAM2 segmentation refinement **(Figure S7D)**. Therefore, we employed two manual segmentation strategies to define the ground truth: labeling using red images **(Figure 2F)** and labeling using gray-scaled versions **(Figure 2G)**. As expected, we observed performance drops when using an SIOU or IOU threshold of 0.5 **(Figures 2H & S8A)**. For example, with SIOU at a default confidence of 0.7, the green H2B-GFP, gray H2B-mCherry, and red H2B-mCherry images have F1 scores of 0.955 (**Figure 2E)**, 0.808, and 0.706 **(Figure 2H)**, respectively. Furthermore, we consistently observed superior performance in the manual labeling of MN using the gray-scaled images. However, by adjusting the IOU threshold to 0.1, thereby prioritizing MN detection over precise segmentation, we were able to improve performance **(Figure S8B)**. In conclusion, we determined that the model was robust against variations in color while reliably detecting and reasonably segmenting MN in low-quality images.

Finally, we compared the performance of our model to micronuclAI for scoring CIN^19^. Briefly, micronuclAI is trained to count both MN and nuclear buds (NBUDs). NBUDs can indicate CIN, but they are not physically separated from the nuclei. Most importantly, NBUDs and MN can have distinct mechanistic origins and may diverge in their downstream consequences. Therefore, since MN and NBUDs are, by definition, unique structures, our comparison aims to demonstrate that training strategies must be considered when making predictions. Upon obtaining expert MN counts from H2B-GFP images (n=60), we found that MN counts from microNucML align more closely with the expert’s manual count (*β*=1.020, ρ=0.801, p-value = 3.597e-28) compared to micronuclAI CIN scores (*β*=1.496, ρ=0.556, p-value = 4.044e-11; **Figure 2I**). Furthermore, we noted that no filtering was required for apoptotic-like segmentations among our model’s predictions.

### Application of microNucML for MN counting

After carefully establishing that microNucML can precisely detect and segment MN, we applied the model to immunofluorescence data obtained from the MCF10A cell line exposed to DNA damage conditions to gain biological insights into genome instability. Specifically, we investigated the downstream consequences of MN formation on innate immune signaling via the cGAS-STING pathway by merging MN segmentations with the colocalization of cGAS. In tandem, we highlighted a novel strategy for characterizing MN production by integrating MN and nucleus segmentation.

MN can only appear following cellular division^6^. Due to variations in cell division rate and cellular seeding, it is standard to normalize MN counts by the number of observable nuclei in an image. Once identified, MN are often classified as “ruptured,” with their DNA now exposed to the cytoplasm, or “intact,” with transport-deficient approximation of a nuclear envelope surrounding their DNA fragment. This classification is relevant to the cGAS-STING signaling pathway, as it can be activated by cytosolic DNA, functioning as part of the immune system, whereby ruptured MN serve as a cytosolic DNA reservoir^6,25,26^. Importantly, not all MN rupture (i.e., no cytosolic DNA access), and not all ruptured MN exhibit cGAS recruitment^7,27^.

Therefore, we utilized the normalized MN counts obtained from H2B-GFP and cGAS-mCherry cells to analyze temporal aspects of MN rupture. Some wells were arrested at the G2-M boundary for 24 hours, after which all cells were exposed to varying doses of irradiation (IR; 2-20 Gy). We began by segmenting MN **(Figure 3A)**, identifying pixels with a strong red (cGAS-mCherry) hue **(Figure 3B)**, then counting cGAS+ MN **(Figure 3C)**. We observed that 24 hours post-IR, increasing IR doses are correlated with enhanced MN production, except at 20 Gy **(Figure 3D)**. However, regardless of dose, the G2-M stalled population generated fewer MN, while displaying a higher proportion of cGAS+ MN. The discrepancy between MN production and cGAS response dissipated 72 hours post-IR (**Figure S9**). This is in alignment with published data using manual counts of cGAS+ MN after IR^6^. These results indicate that MN production, followed by cGAS response, may depend on cell cycle progression, be time-sensitive, and could be influenced by other factors, such as nuclear envelope fragility or MN content^28,29^. Overall, we found that the tool could quickly compute MN counts and seamlessly integrate with a colocalization study.

**Figure 3.**
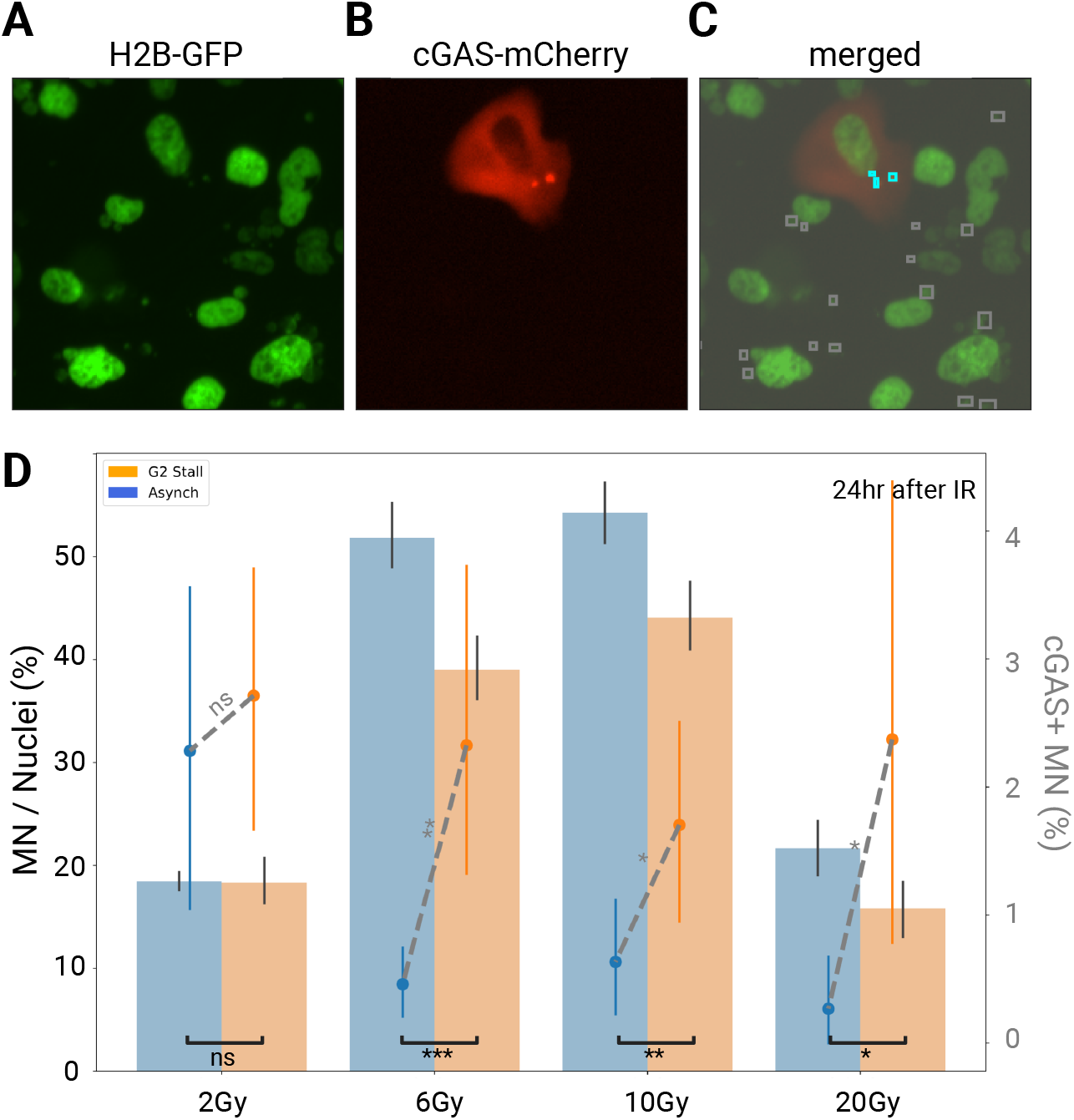
Application of MicroNucML-based segmentation of MN with cGAS-mCherry localization. Among cell constructs with H2B-GFP **(A)** and cGAS-mCherry **(B)**, both histones and cGAS responses were visualized and integrated. Upon MN segmentation with MicroNucML, MN were discriminated by the presence and absence of cGAS recruitment **(C)**. 24 hours after irradiation (IR), at varying doses, some cells underwent G2 stalling. **(D)** The barplot, which aligns with the left axis and black statistical results, demonstrates the MN counts relative to the nuclei counts. The point plot, which depicts the proportion of cGAS+ MN, is aligned with the right axis and accompanied by gray statistical results (Mann-Whitney U test). Asynchronized and G2 stalled cell populations are in blue and orange, respectively. Error bars show the 95% confidence intervals.

### Application of microNucML to refine MN and parent nuclei relationships

Although normalizing MN to the nuclei population serves as a proxy for CIN, not all nuclei form MN. MicroNucML establishes distance-based nuclei-MN relationships, which can be used to calculate the number of MN per parent nucleus and to assess the relative size of MN. To demonstrate these features, we began with cells that underwent transient G1 stalling and/or ATR inhibition (ATRi), which were imaged over a 72-hour period during which there is ample time for mitotic opportunities to produce MN. ATRi induces replication stress by permitting dormant replication origin firing and slowing replication forks, in part by depleting nucleotide pools^30^. Furthermore, ATRi has been directly associated with CIN^31^ and MN formation^32^. Therefore, following the release of G1 stalling, as these cells enter the S phase, we expected them to exhibit greater and more immediate sensitivity to ATRi by producing more MN after their next mitosis.

We applied a linear mixed model using normalized MN counts and observed that both G1 stalling and ATRi significantly increase MN production (*β*_Stall_=-0.054, p-value_Stall_=0.003, *β*_ATRi_=−0.160, p-value_Stall_=0.0, **Figure 4A**). Interestingly, we found a 13-hour delay in peak normalized MN counts between the ATRi populations, which were left asynchronized, and those that were stalled. We repeated this analysis using MN per parent nuclei, which served as a proxy for the number of DNA fragments originating from either unresolved breaks or mitotic errors. In this case, we found that ATRi alone impacts MN per parent nuclei (*β*_Stall_=0.011, p-value_Stall_=0.613, *β*_ATRi_=−0.137, p-value_Stall_=0.0; **Figure 4B**). Subsequently, we used the pixel area of a parent nucleus and its MN to measure relative MN size. In H2B-GFP constructs, the relative size of MN may depend on the amount of severed DNA but may be skewed by factors such as DNA compaction or cell cycle state, although we expect MN to range from 1/100th to 1/10th the size of the parent nucleus^33^. We found that, regardless of condition or time point, MN generally remained within the expected size range **(Figure S10)**. However, the asynchronous ATRi population consistently produced slightly larger MN (approximately 6/50^th^ the size, between 15-30 hours post-IR). These findings demonstrate MicroNucML’s applicability in analyzing MN dynamics, suggesting that G1 stalling and/or ATR inhibition may influence the speed of MN formation and might act as modulators of the magnitude of instability endured by the cells.

**Figure 4.**
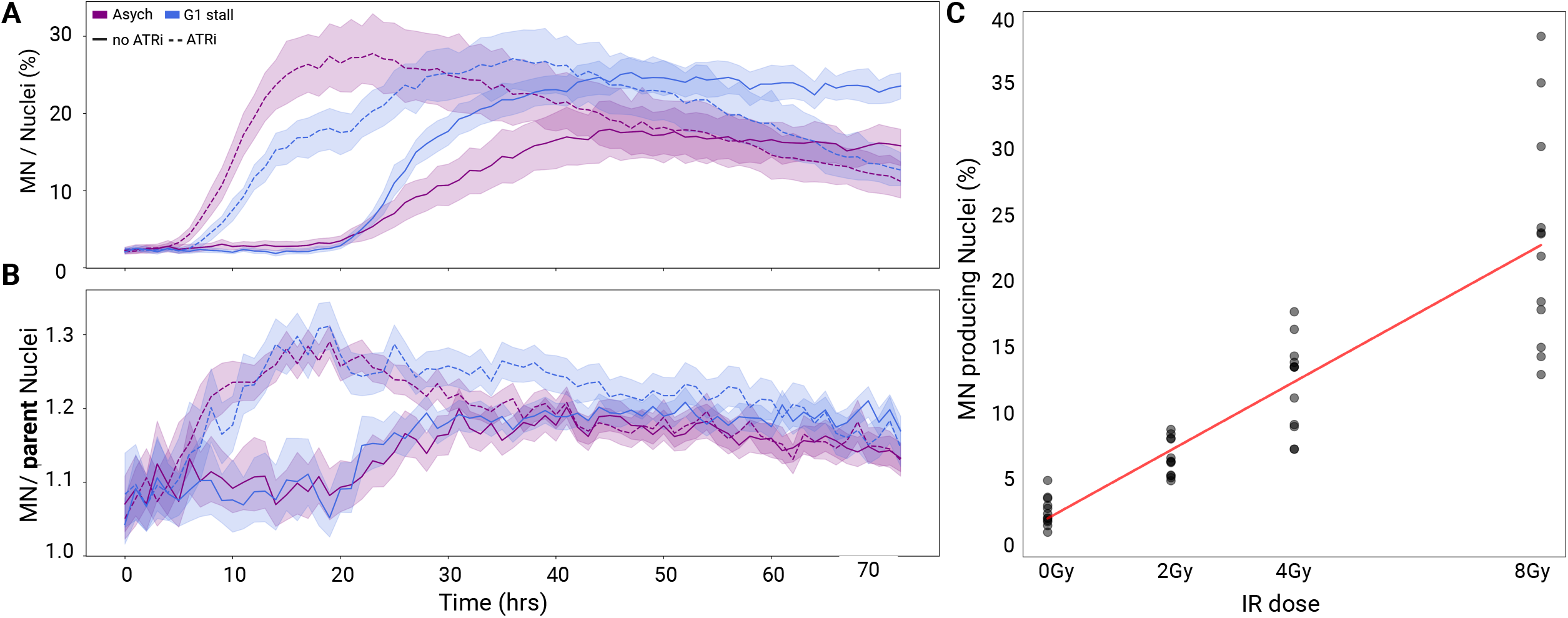
Integration of MN and nuclei segmentation for biological interpretations of MN production. A time-course experiment, following conditions of IR, ATR inhibition, and/or G1 stalling, depicts the mean normalized MN following IR across the four conditions. Shading shows the 95% confidence interval for a given time point **(A)**. Similarly, the normalized MN per parent nuclei are displayed **(B)**, with a minimum of 1 MN per parent nucleus. Asynchronized and G1 stalled cell populations are in purple and blue, respectively. Cells without and with ATRi exposure are shown with solid and dotted lines, respectively. **(C)** Next, a scatterplot highlights the proportion of MN+ nuclei in a population, relative to increasing IR dose. The linear regression is shown in red.

Finally, we aimed to model MN-producing nuclei as a function of increasing DNA damage. Experimental studies utilizing normalized MN counts assume that all nuclei contribute equally to MN production. However, following DNA damage, not all cells maintain the capacity to divide. Some cells become senescent, and others may trigger apoptosis^34^. With our tool, upon identifying parent nuclei, we can more accurately estimate the proportion of nuclei in a population that exhibit MN. We found that 48 hours post-IR, one radiation unit contributes to a 2.6% increase in MN+ nuclei (*β*=2.63, ρ=0.8701, p-value = 9.70e-16; **Figure 4C**). Therefore, we propose that automated nuclei and MN counting by MicroNucML, along with resolving their spatial relationships, can assist in refining dose-response models at scale.

## Discussion

Exploiting MN as a measure of genotoxicity has been a long-standing effort. However, the traditional manual approach to segmenting and counting MN can be labor-intensive. Additionally, thresholding strategies are susceptible to experiment-specific variation. In this work, we developed a robust ML model and an associated tool for high-throughput counting and segmentation of MN and nuclei. Our tool is available as an open-source Python package with minimal dependencies and low runtime, requiring only basic image preprocessing supported by a GPU.

We employed a two-phase manual labeling strategy to build our model by reducing label burden and enhancing segmentation quality. In the first phase, we developed a binary classifier that streamlined the population of tiles requiring labeling. Consequently, in the second phase, we established the largest publicly available repository of labeled MN segmentations. Utilizing LabelStudio’s brush tool, we corrected for overdrawing by incorporating the user-drawn masks as prompts for SAM2. Upon visual inspection, we observed that the SAM2 foundation model tightens the masks, thus improving the quality of the ground truth. This strategy aligns with previous work, in which SAM2 uses prompts to effectively segment underwater objects^35^. Furthermore, the brush-generated masks act as prompts to help SAM2 concentrate on the regions of interest. By adopting this approach, we effectively transferred the high-level knowledge embedded in SAM2 into our training dataset, without needing to retrain the entire model. While SAM2 is mainly utilized for auto-labeling in our workflow, this prompt-guided refinement parallels concepts in knowledge distillation^36^, whereby a powerful “teacher” model (i.e., SAM2) provides enhanced supervision for training a “student” model (i.e., our MN segmentation model). This, therefore, can mitigate variation across labelers, a common issue that affects biological tasks^37^.

Subsequently, during post-processing, the tool evaluates the density of MN to filter out cells exhibiting blebbing or apoptosis. Since apoptotic cells were not labeled during training, as expected, this filter was not triggered when applied to an independent set of images. However, we aimed to strengthen the tool to make predictions within biological expectations, specifically regarding the density of MN, which adheres to the expected MN burden^38^. We suggest that such strategies can function as “prediction safeguards”, which can be adjusted or disabled by the user.

At the end of the workflow, the tool applies a distance-based assignment between MN and their parent nuclei. These assignments remain agnostic to the cell’s behavior throughout division. We propose that accurate assignments could be achieved through cell tracking during division or by inclusion of a plasma membrane dye to delineate cellular borders. Previous groups have attempted to develop and evaluate cell tracking systems; however, this task remains challenging^39,40^. Furthermore, these models overlook evidence of CIN. Developing such a model would require capturing images at an appropriate frequency, along with classification and segmentation across frames.

We trained and evaluated several MN segmentation models, achieving state-of-the-art performance with an optimized Mask-RCNN model, which we named MicroNucML. Interestingly, when comparing Mask-RCNN models to those with YOLO architectures, we found that the former outperformed the latter unexpectedly. While YOLO excels at general tasks^41^, earlier studies have concluded that Mask-RCNN is robust, especially for segmenting small and variable objects^42^. Furthermore, in biological contexts, Mask-RCNN has demonstrated superiority over YOLO^43^. This suggests that the optimizations we implemented, such as adjusting anchors and IOU, likely contributed to these performance differences.

In a similar fashion, we assessed the quality of our model’s segmentations and our evaluation metric, IOU, by considering the size of MN. We found that MicroNucML can appropriately predict MN size and position. However, the canonical IOU is sensitive to small errors, which we attributed to the small size of MN. We alternatively used SIOU, which aims to assess the segmentation of small objects while aligning with human perception of accuracy^24^. Although SIOU has yet to be widely deployed for biological tasks, our benchmarking suggests that when pursuing small object detection, IOU should be used in conjunction with alternative evaluation metrics^44^. Importantly, accurate predictions of MN size are biologically relevant, since MN size can be an indicator of properties of the enclosed DNA, along with the downstream risk of envelope rupture^45^. For example, MN containing highly compacted chromatin will be smaller compared to MN with accessible chromatin, further implicating proteins that may later bind to the DNA, such as cGAS^46–49^.

Next, we employed data augmentation strategies primarily aimed at addressing color and image quality to enhance our model’s generalizability for MN detection. We manipulated the color of the held-out H2B-GFP images, observing consistency in the model’s performance across distinct colors. Consequently, this suggests that our model can handle images of different visualization strategies (green, red, blue via GFP, mCherry, DAPI, Hoechst dyes) while maintaining very high segmentation quality. However, when it comes to images with markedly poorer quality, the model appears to struggle slightly with precise segmentations, although it can still produce accurate MN counts. Despite this, the small decrease in performance was expected, since it is well established that the performance of ML models trained on microscopy images can vary due to variations in image quality^37^. Nevertheless, ML models have been shown to outperform threshold-based strategies, particularly when identifying an object’s contour^37^. Thus, with respect to poorer image quality, our model likely remains more effective compared to thresholding.

Furthermore, we benchmarked our model against another existing tool, micronuclAI^19^. Compared to our approach, micronuclAI counts MN and NBUDs. Both MN and NBUDs can serve as markers for genomic instability^50,51^, with some MN being established through a prolonged budding period^52,53^. However, their origins and downstream fates can differ. For example, NBUDs, which commonly originate from interstitial fragments^20,21^, and by definition, do not possess an independent nuclear envelope (i.e., relevant for cGAS response)^54^. We utilized an external MN dataset for benchmarking and observed that our model aligned more closely with expert MN counts, as anticipated, since our expert excluded NBUDs. Additionally, the underlying model architecture differs between the two models. For instance, micronuclAI is trained as a regression task, while MicroNucML conducts segmentation. Previous studies suggest that segmentation tasks commonly outperform regression due to increased task complexity^55,56^. Furthermore, since their regression task exclusively provides a CIN score instead of masks, users are limited in their ability to quality control MN or NBUD identification. Therefore, we recommend that users consider their specific questions and applications when choosing a model for MN detection.

After establishing the robustness and generalizability of micronucML, we applied our method to images specifically taken to explore MN production in various contexts of genomic instability. For instance, an inspection of MN from a H2B-GFP and cGAS-mCherry colocalization study revealed variations in MN production and the strength of cGAS response. Generally, this aligns with previous work characterizing the modulation of inflammatory responses following DNA damage as time- and cell cycle-dependent^28,29^. Recent research has explored other features, such as histone modifications, that discriminate between MN with and without cGAS recruitment^27^. Therefore, future higher-throughput multiplexed experiments, which segment MN using MicroNucML, could identify a broader range of features that differentiate MN fate.

Subsequently, we explored ways in which MicroNucML could be utilized to refine quantifications of MN production. Importantly, MN arise after a cell completes division. First, we aimed to investigate whether G1 stalling and/or the replication stressor ATRi alter the speed or burden of MN production. Classically, MN production is measured by normalizing MN to the nuclei population. Using this technique, we observed that both G1 stalling and ATRi modify MN load and production speed following IR. However, not all nuclei produce MN, and some nuclei may produce multiple MN. Through MN-parent nuclei assignment, we further clarified that cells subjected to ATRi uniquely produce more MN per parent nucleus, along with slightly larger MN. This can serve as a proxy for DNA fragmentation or the number of lagging chromosomes^57,58^. Furthermore, MN size is known to vary depending on the DNA-damaging agent^59^. Although the entire collection of ATRi effects is still under investigation, we propose that replication stress itself and/or weakened nuclear envelopes could explain the observed increase in MN per cell^58,60,61^. Future research could leverage the provided collection of MN metrics as a strategy to investigate mechanisms of DNA damage, while also differentiating between parent and non-parent nuclei.

Overall, our study provides a comprehensive description of the training and deployment of an accurate MN segmentation model and tool. Despite its accurate performance, our current model has areas for improvement. In particular, the training dataset used to construct our model remains largely homogeneous. For example, we trained the model using experiments from two human cell lines (i.e., MCF10A and RPE-1) with H2B-GFP constructs, which were imaged using a single instrument (i.e., Incucyte) at a single magnification (i.e., 20x). In the case of external datasets, which require preprocessing of images to match the magnification of the training images, we expect degradations in MN segmentation quality due to losses in image resolution. Furthermore, our predictions focused on MN, paired with automated nuclei counting. Therefore, dividing and apoptotic nuclei states were excluded from our analyses. However, for studies aimed at exploring the complete landscape of DNA damage, greater granularity among cell states may offer deeper biological insights. More simply, segmentation efforts could incorporate multiclass nuclei labels, including specific phases of the cell cycle, or, as previously mentioned, integrate cell tracking. This may be beneficial for comparing cell populations that display senescence, remain actively dividing, or undergo death as a strategy to differentiate the consequences of CIN. Despite these limitations, our method offers high accuracy, is computationally efficient, and can be widely applied across various experimental studies. Overall, our tool will be an effective method for characterizing MN production, thereby establishing its role in genomic instability.

## Methods

### Description of images used for manual labelling and training

Our labeled training data consisted of images captured by an Incucyte, a live image microscope, at 20x magnification and a 0.625 µm/pixel resolution. We obtained training images from H2B-GFP constructs in MCF10A and RPE-1 cell lines. Our goal was to capture a variety of nuclei and MN sizes, morphologies, and densities. Therefore, we labeled and trained models under various experimental conditions. For example, cells were exposed to different genotoxic stressors, including irradiation and/or drugs such as HU, DRB, and MMS, along with ATR inhibition, G1 or G2 cell cycle stalling, gene knockouts (KO; CRISPR screens or p53 KO constructs), and variations in the timing of image captured post-stressor, which ranged from hours to days. Cells were subjected to radiation using a Gammacell 40 extractor, which transmits gamma radiation.

Manual labeling was generally conducted by three experts in two phases using LabelStudio^22^. First, we employed a polygon tool to segment nuclei and MN from full-sized images (1400×1000 pixels). We utilized these images (n=148) to develop a binary classifier for identifying regions of interest with a high probability of MN. After creating smaller image tiles (224×224 pixels), we applied the binary classifier to filter tiles containing MN, thereby reducing the labeling burden. The subset of tiles was then used to segment MN with the brush tool. These images were used to train the final segmentation model.

### Training a binary classifier to identify regions of interest

Regardless of the experimental conditions, only a small proportion of cells produce MN. This presents a practical challenge for both manual and automated segmentation, as it requires identifying small objects within a large search space. To address this challenge, full-sized images with segmented MN (using the polygon tool) were employed to train a convolutional neural network (CNN) for classifying tiles based on the presence or absence of MN. The image set consisted of 967 MN+ tiles and 1,382 MN-tiles. We utilized a ResNet101 model pretrained on ImageNet, adding two fully connected layers. We conducted five-fold cross-validation (CV), using 90% for training and 10% for validation. Further, we applied data augmentation by randomly flipping tiles during training. We assessed the model using average accuracy and then produced a confusion matrix by applying the model across the entire image set.

### Building a model for MN segmentation

After applying the binary classifier to unseen images, we generated a subset of tiles with a high probability of MN presence. These tiles were then provided to the experts, who segmented MN using LabelStudio’s brush tool with a single click. This strategy reduced the search space, resulting in a more efficient labeling process by increasing segmentation speed and alleviating the labeling burden. Next, since brush tools often overdraw an object’s boundaries, we refined the MN segmentation. We used the original MN masks as input prompts for SAM2, a foundation model, to tighten the segmentations. Consequently, the refined segmentations served as ground truth for model training. Afterward, we partitioned the tiles into a training dataset (80%) and a testing dataset (20%), employing cross-validation during training. This resulted in a training dataset of 1,529 tiles with 2,967 refined MN, and a testing dataset of 300 tiles with 614 refined MN, which jointly come from 687 full-sized images.

Our goal was to develop a model that is invariant to color and image quality, enabling users to deploy it with various MN visualization strategies (i.e., gray-scaled, red, blue, green, etc.) on images with differing background fluorescence from multiple instruments. However, our training images consisted exclusively of Incucyte images from H2B-GFP constructs. To address this homogeneity, we employed a series of data augmentation strategies. For color, we randomized the images’ brightness, contrast, saturation, and hue within a ±30% range. We applied Gaussian blurring with a kernel size of 5×9 and a standard deviation ranging from 0.1 to 2 for image quality. Finally, we incorporated random horizontal and vertical flips with a probability of 0.5.

Previous studies that performed MN detection predominantly employed CNN-based approaches. To address the challenges of segmenting small and highly variable objects, we explored You Only Look Once (YOLO) and Region-based CNN (RCNN) architectures with different backbones. These two architectures utilize single-stage and two-stage object detection frameworks, respectively. The Mask RCNN model includes a stage for region proposal, followed by object classification. We fine-tuned several backbone networks, including ResNet-50, Swin Transformer, and ResNeXt101. The backbones are pre-trained on the ImageNet-1K dataset. We rationalized that this was appropriate since backbones trained with general image sets, such as ImageNet-1K, can be applied to specific datasets upon fine-tuning. Thus, we included batch normalization, added convolutional layers, and froze all but the last three layers of the network to allow for this domain-specific adaptation. To enhance feature extraction across scales, we applied a Feature Pyramid Network (FPN) to the five-level feature maps generated by the backbones. These maps are then passed to region proposal networks (RPNs) to identify candidate regions. Among the RPNs, we varied the size and quality of anchors. Given that MN are small and can vary in morphology, we tested a variety of anchor sizes (4, 8, 16, 32, 64, 128) and two intersection over union (IOU) thresholds (0.5 and 0.7). To evaluate our optimization attempts, we performed ablation studies with models trained for up to 10 epochs.

We fine-tuned our model using the AdamW optimizer because of its adaptive learning rate and efficiency in managing sparse gradients. The learning rate was set to 0.001, with momentum values of 0.9 and 0.999 for the first and second iterations, respectively. We closely monitored the training progress to ensure optimal convergence, implementing early stopping based on validation performance to avoid overfitting. Optimization was conducted with loss functions, including for classification (weighted categorical cross entropy), box regression (L1), and mask (cross entropy), to balance segmentation accuracy and generalization. To eliminate duplicate predictions from the model, we applied the non-maximum suppression algorithm with an IOU threshold of 0.2. We evaluated the model’s performance using precision, recall, mean average precision (mAP) of MN segmented with a minimum IOU of 0.5 (mAP@50), and F1 score, requiring segmentations to achieve a minimum confidence of 0.7.

Importantly, subject to explorations in prediction quality, we evaluated segmentations using the IOU, along with the scale-adaptive IOU (SIOU). SIOU has been described as aligning with human perception of accuracy and emphasizes both localization accuracy and object size^24^, thereby making it useful when conducting small object detection. SIOU is described as;

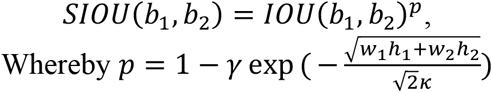

Whereby, b_1_ and b_2_ represent the location of the ground truth and predicted boundary boxes, respectively, while p represents the scaling factor, as a function of the object sizes. In this case, we fixed γ = 0.5 and κ = 64 as proposed by an earlier study^24^. Finally, we measured the performance trade-off for varying segmentation confidence by comparing F1 scores with confidences between 0.1 and 0.9 with a step size of 0.1. Finally, we monitored inference time using two different GPUs: T4 and A100.

### Post-processing MN segmentation by filtering and nuclei assignment

After genotoxic stress, some cells produce MN, while others may undergo apoptosis. Nuclei that are preparing for or undergoing apoptosis may appear fragmented or “blebbed”; visually, this can resemble a small area with a high density of MN. Therefore, we employed Density-Based Spatial Clustering of Applications with Noise (DBSCAN)^62^ to filter out MN in these high-density areas. Using a maximum neighbor distance of 20, we eliminated MN clusters containing five or more MN.

In high-throughput studies, assigning MN to their parent nuclei can be challenging. We aimed to automate these assignments using a distance-based approach. Notably, this strategy disregards the behavior of proposed parent nuclei during division and operates on the presented image (“memoryless”). This limitation was deemed acceptable, as enhancing accuracy would require tracking specific nuclei across time and cell cycle states, which is challenging due to image variability between the imaged timeframes. First, alongside MN segmentation, we segmented all nuclei in the image using SAM2. We rationalized this approach since SAM has been previously shown to be an effective cell segmentation strategy^63^. We used default settings, except for the ‘point per side’ parameter, which we adjusted to 128 to account for the density and size of the nuclei. Since SAM2 is agnostic to MN segmentation, it can, in theory, segment larger MN. To address these instances, we used the image’s maximum MN size as a threshold when filtering out segmented nuclei. While SAM2 is employed in our downstream analysis, our tool adapts a framework that accepts any nuclei segmentation masks, enabling flexibility when integrating with MN segmentation. Next, we identified the centroids of all the MN and nuclei in the image. For each MN, we identified the three closest nuclei centroids. These nuclei are then segmented into a closed polygon, from which we calculate the Euclidean distance from the centroid to a line segment. Thus, we denote the parent nuclei as the nuclei with the smallest distance between the MN centroid and the nearest edge of a nuclei.

### Improving model generalizability by color and image quality invariance studies

As described, we applied a series of data augmentation strategies to address the homogeneity in color presented by the green (H2B-GFP) images used during training. Therefore, we aimed to investigate the model’s robustness using images that mimic other DNA visualization techniques (i.e., gray-scaled, red, blue from mCherry, DAPI, or Hoechst) while maintaining similar image quality. We converted the green test set of images into gray, red, or blue by adjusting the hue of the images by ±60° after converting them from RGB to HSV format. We evaluated the model’s performance across the four sets of images using the F1 score (at an IOU and SIOU of 0.5) and employed segmentation confidence thresholds ranging from 0 to 1.

Next, we aimed to evaluate the model’s generalizability by using H2B-mCherry images that were unseen by the model and outside the distribution of the training images. In this case, we had 71 full-sized images, which produced 308 tiles with 477 MN. Most notably, this set of images fluoresces H2B in red and appears to be of poorer quality. To confirm, we gray-scaled the test set images (i.e., H2B-GFP) and the H2B-mCherry images, then compared the MN size and variation in pixel value from edge-filtered images, which represents image sharpness. Importantly, poorer quality presented a challenge when manually segmenting H2B-mCherry images. We noted that nuclei and MN are more pronounced with gray scaling. Therefore, we implemented a blind double-labeling approach when producing manual segmentations. Two independent labelers annotated the same set of H2B-mCherry tiles using the LabelStudio brush tool, utilizing either the red or gray-scaled set of images. To maintain consistency, we refined the MN masks using SAM2, thereby creating two sets of ground truth. Subsequently, we applied MicroNucML. During evaluation, we aimed to measure the influence of the manual segmentation strategy and image quality. Therefore, we calculated F1 scores across varying confidence scores using IOU and SIOU thresholds at 0.5 and 0.1, thereby prioritizing segmentation quality and MN counting, respectively.

### Comparison with micronuclAI

We compared our method to another CIN detection method, micronuclAI. MicronuclAI is trained to predict a CIN score using both MN and nuclear buds (NBUDs). It employs these counts to perform a regression task by utilizing the area surrounding previously segmented nuclei. For this study, we manually counted MN in a held-out dataset of MCF10A cells imaged using the Incucyte (n = 60). Simultaneously, we predicted MN counts for the images using our model, MicroNucML, as well as CIN scores from micronuclAI. To generate counts with micronuclAI, we applied the default settings and used Mesmer to segment the nuclei, aiming to replicate their workflow. We evaluated and compared model performances by creating linear regressions between expert counts and predicted counts.

### Application of microNucML for downstream analyses

After training and testing the model, we aimed to demonstrate use cases that highlight the biological insights obtained from the tool. To begin, we conducted a series of experiments to showcase its integration with colocalization, replication stress, and dosimetry studies. The bottoms of all the images were cropped (1400×950 pixels) to remove text. Additionally, we utilized MCF10A cells with H2B-GFP constructs. We applied the MicroNucML model using the default confidence threshold of 0.7.

First, the colocalization study involved cells containing both H2B-GFP and cGAS-mCherry constructs. Additionally, some cells were treated with the CDK1 inhibitor RO-3306 at a concentration of 10 µM for 24 hours. Next, all cells were exposed to radiation at varying doses (2, 6, 10, and 20 Gy). The cells were then imaged one, two, and three days post-radiation (n = 288). We applied the MicroNucML model and concurrently analyzed the mCherry-cGAS images independently. We identified red pixels by utilizing the 99th percentile of red intensity across all images. The sufficiently red pixels were overlaid with MN segmentation, allowing us to calculate the proportion of MN displaying a cGAS response.

The replication stress study aimed to demonstrate the interplay between replication stress, induced by an ATR inhibitor (ATRi; VE-821 at 2.5μM), and cell cycle stalling at the G1 phase. First, some cells were starved for 24 hours to induce G1 stalling. Next, some cells were subjected to ATRi for 1 hour. Subsequently, all cells were irradiated with 10 Gy and imaged every hour for 72 hours (n = 3,504 full-sized images). To investigate the relationship between nuclei and MN, we calculated the ratio of MN to nuclei, identified the number of MN per parent nucleus, and determined the relative size of MN compared to the parent nuclei. We modeled the dynamics of MN over time using linear mixed models. The wells represented random effects due to seeding variations, while time, G1 stall status, and ATR inhibition served as fixed effects.

For the dosimetry study, cells were subjected to varying radiation doses (0, 2, 4, 8 Gy). Four days after IR, cells were imaged (n = 48), and we employed linear regression to explore the relationship between dose and the proportion of parent nuclei in the population.

## Source code and data availability

The dataset of images and refined SAM2 masks used for training and evaluation is available through Zenodo (https://zenodo.org/records/15312291). The binary tile classifier and optimized Mask-RCNN model can be accessed through HuggingFace (https://huggingface.co/ccglab22/mnClassifier; https://huggingface.co/kew1046/MaskRCNN-resnet50FPN). The open-source Python package, which includes the deployment of MicroNucML and postprocessing parameters, can be found on GitHub (https://github.com/kumarlab-compomics/MicroNuclei_Detection).

## Acknowledgments

SK and NB acknowledge support from the Princess Margaret Cancer Foundation, Canada Research Chair Program, and Terry Fox Research Institute. SH acknowledges support from the Canadian Institute of Health Research and Terry Fox Research Institute.

