## Supplementary Figures for "MicroNucML: A machine learning approach for micronuclei segmentation and the refinement of nuclei-micronuclei relationships"

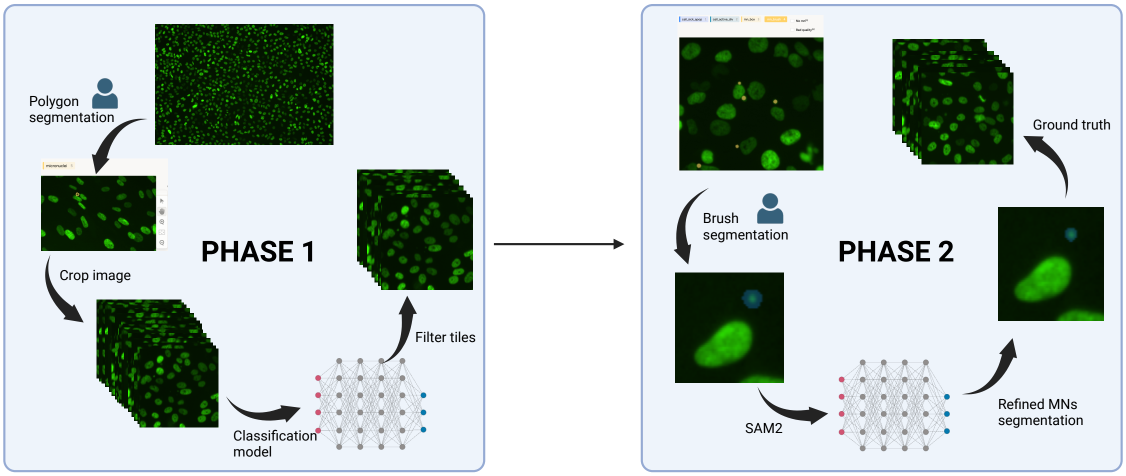


**Figure S1. Two phase approach, designed to increase manual label output.** In phase 1, labellers were presented with full sized images. They were tasked with polygon segmentation of MN using LabelStudio. Smaller tiles were generated, whereby MN+ and MN- tiles were used to train a binary classifier. This classifier was used to generate a subset of tiles, which were then assigned to labellers for brush segmentation, in phase 2. Overdrawn MN, were refined with SAM2, producing higher quality MN segmentation, representative of ground truth.

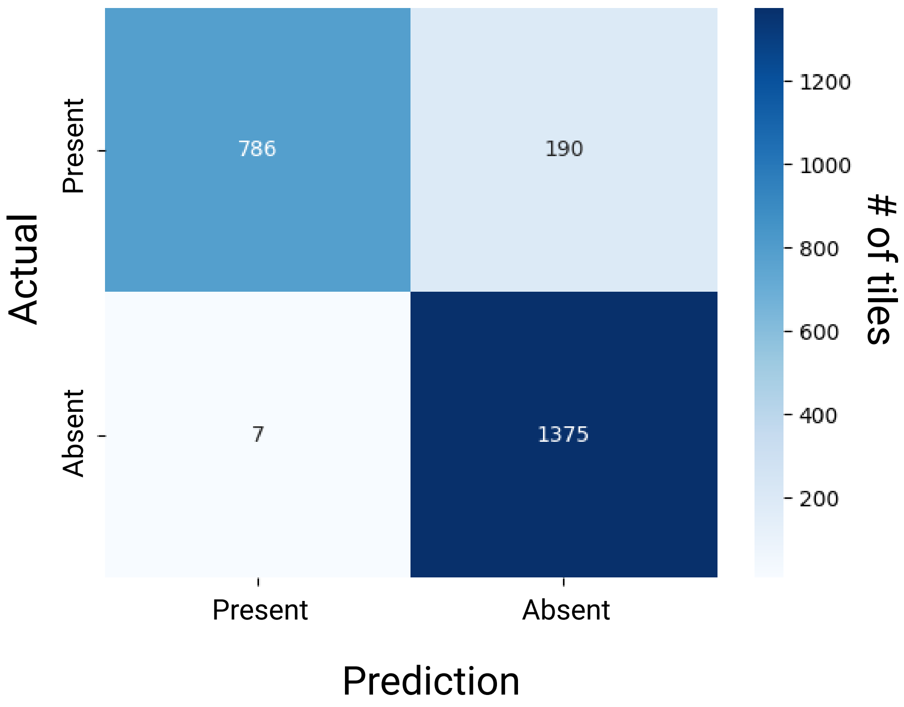


**Figure S2. Confusion matrix produced, as per the binary classifier.** Cases defined as present indicate a tile contains MN, while cases defined as absent, lack MN.

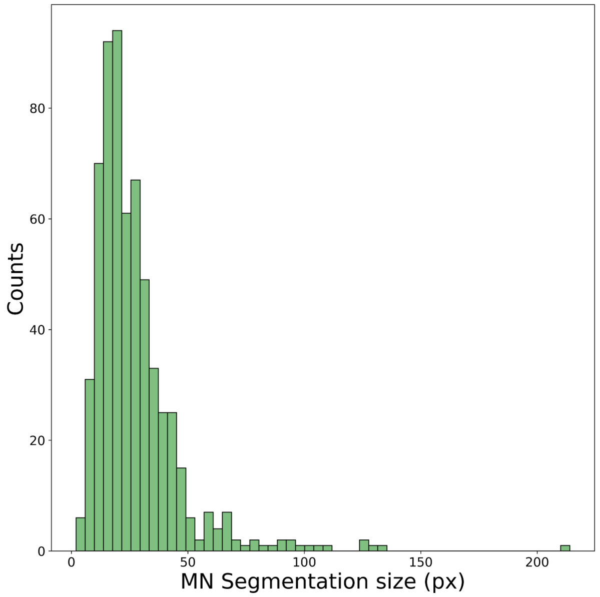


**Figure S3. Histogram of MN sizes (in pixels) in the testing dataset, following SAM2 refinement.**

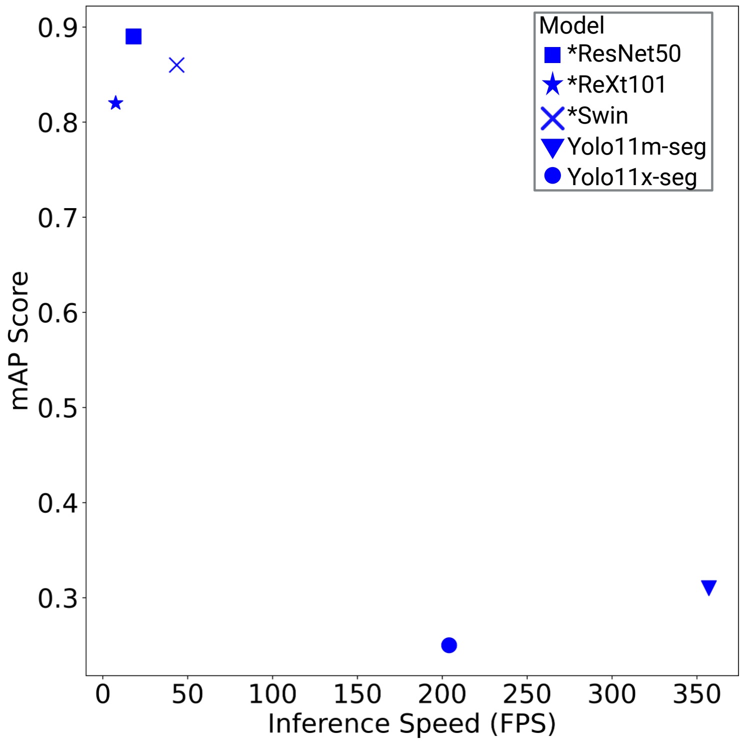


**Figure S4. Scatterplot depicting the inference speed (in frames per second; FPS) compared to the mean average precision (mAP) score across five contending segmentation models.**

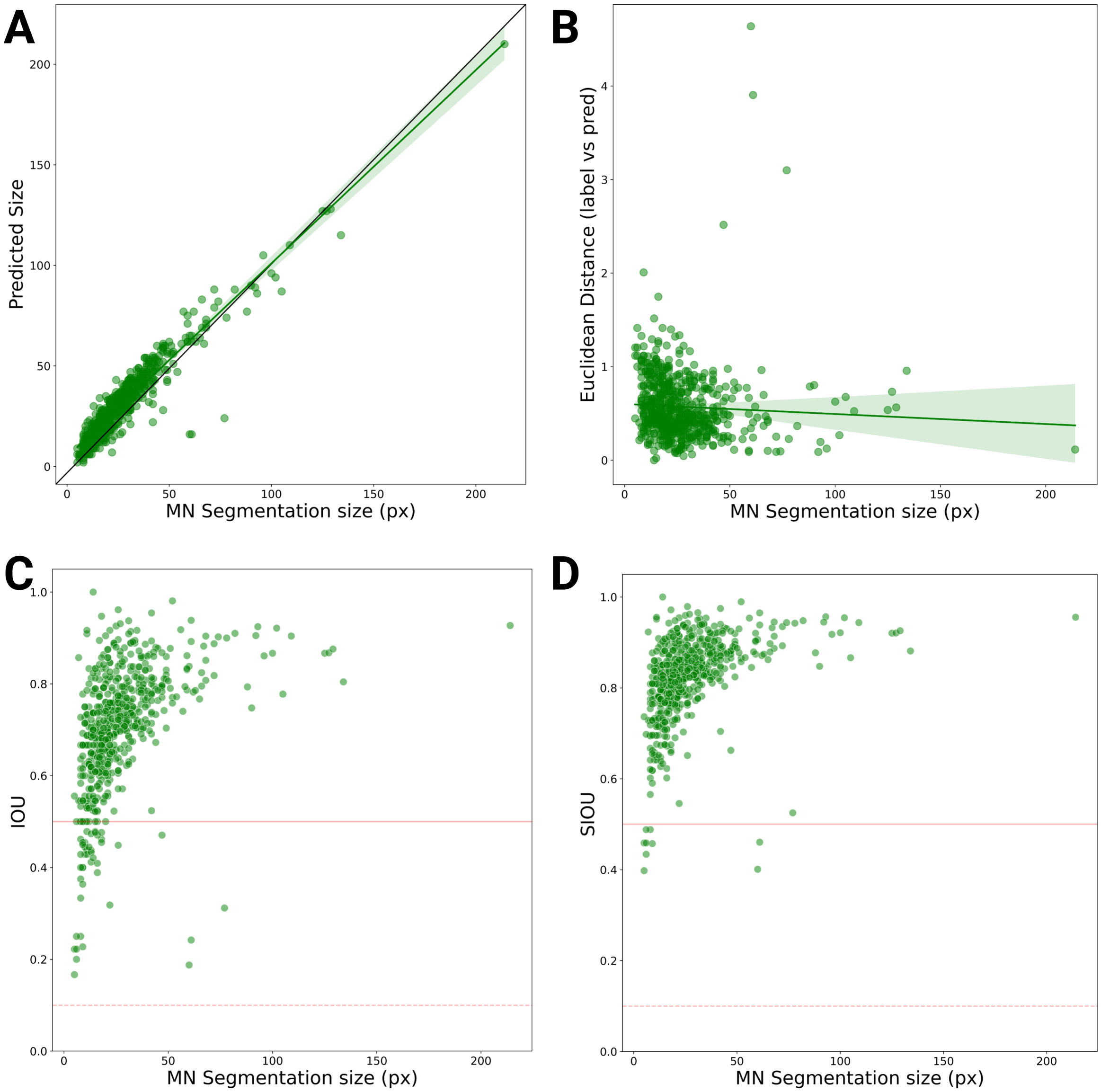


**Figure S5. Rationalization of SIOU and IOU evaluation, in tandem.** (A-B) Exploration into sources of segmentation error. (A) A scatter plot depicting the refined MN segmentation size compared to the predicted size, as per MicroNucML, from the held-out testing dataset. Y=x is in black. (B) A scatterplot comparing MN size to the Euclidean distance, measured between the segmented and predicted position. MN size is in pixels. The linear regressions are in green, with shading as per the 95% confidence interval. (C-D) Comparison between the refined MN size, with IOU (C) and SIOU (D). A horizontal solid and dotted line, depict evaluations at 0.5 (ie. sufficiently well segmented), and 0.1 (ie. poorly segmented, however detected), respectively.

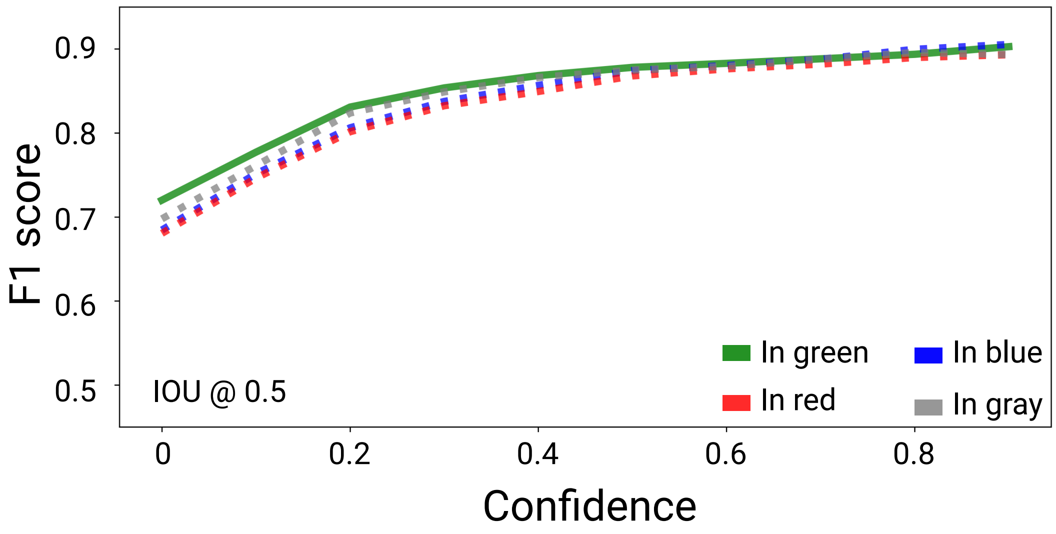


**Figure S6. A line plot depicting colour invariance, as per the H2B-GFP images in green colour, along with red, blue and gray.** As such, upon fixing IOU to a threshold of 0.5, MN confidence and F1 scores were captured across the four colour conditions.

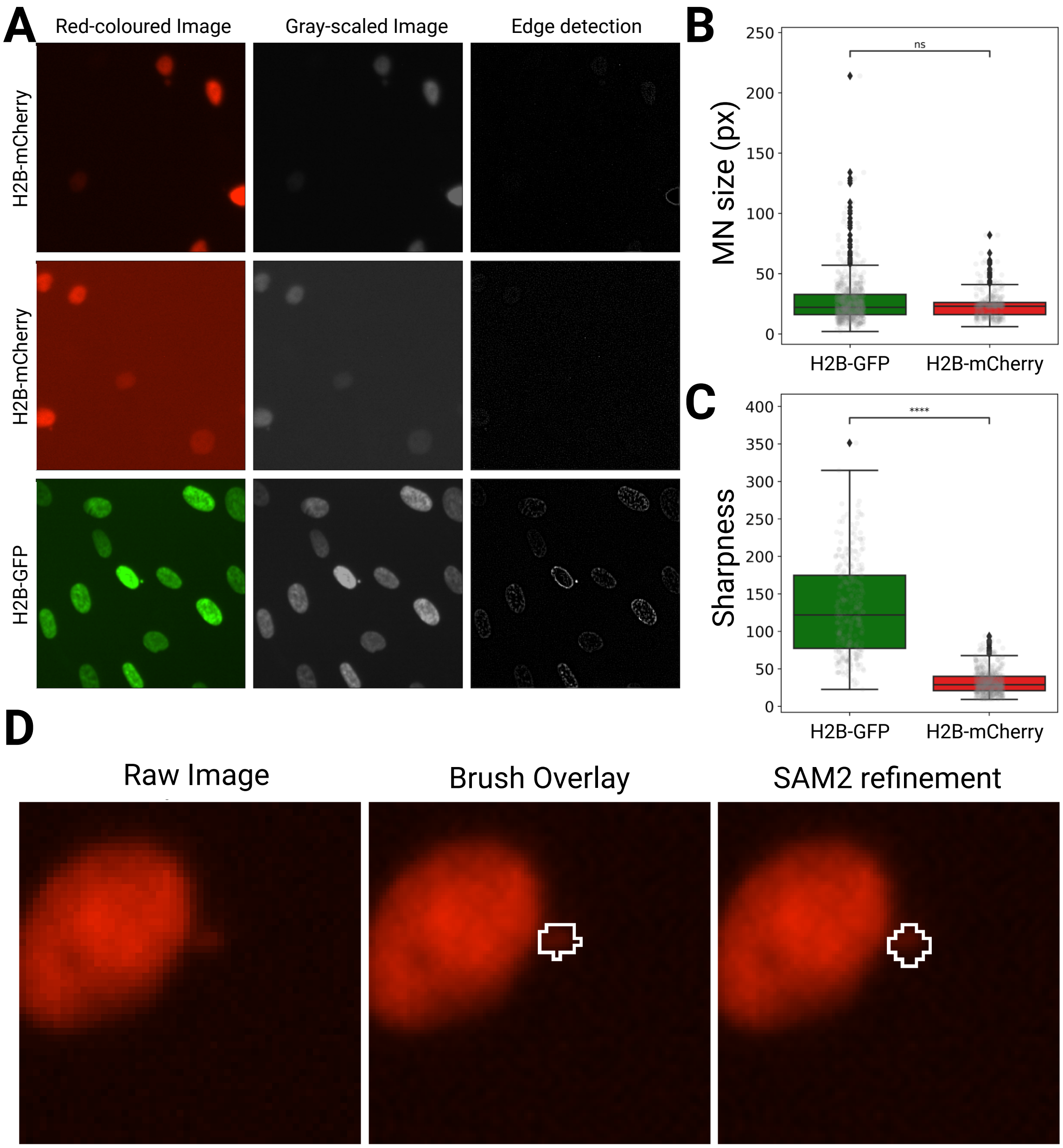


**Figure S7. H2B-mCherry images are of notably poorer quality.** Upon visual inspection, H2B-mCherry images appear out of focus, and have greater background fluorescence, compared to H2B-GFP images (A). Across three tiles, images were gray scaled, then edges of objects were identified with watershedding. Subsequently, MN refined sizes (B), and image sharpness (C) was compared between the H2B-mCherry and H2B-GFP images. Statistical testing with Mann-Whitney U test. Across H2B-mCherry images SAM2 provided minimal improvement (D).

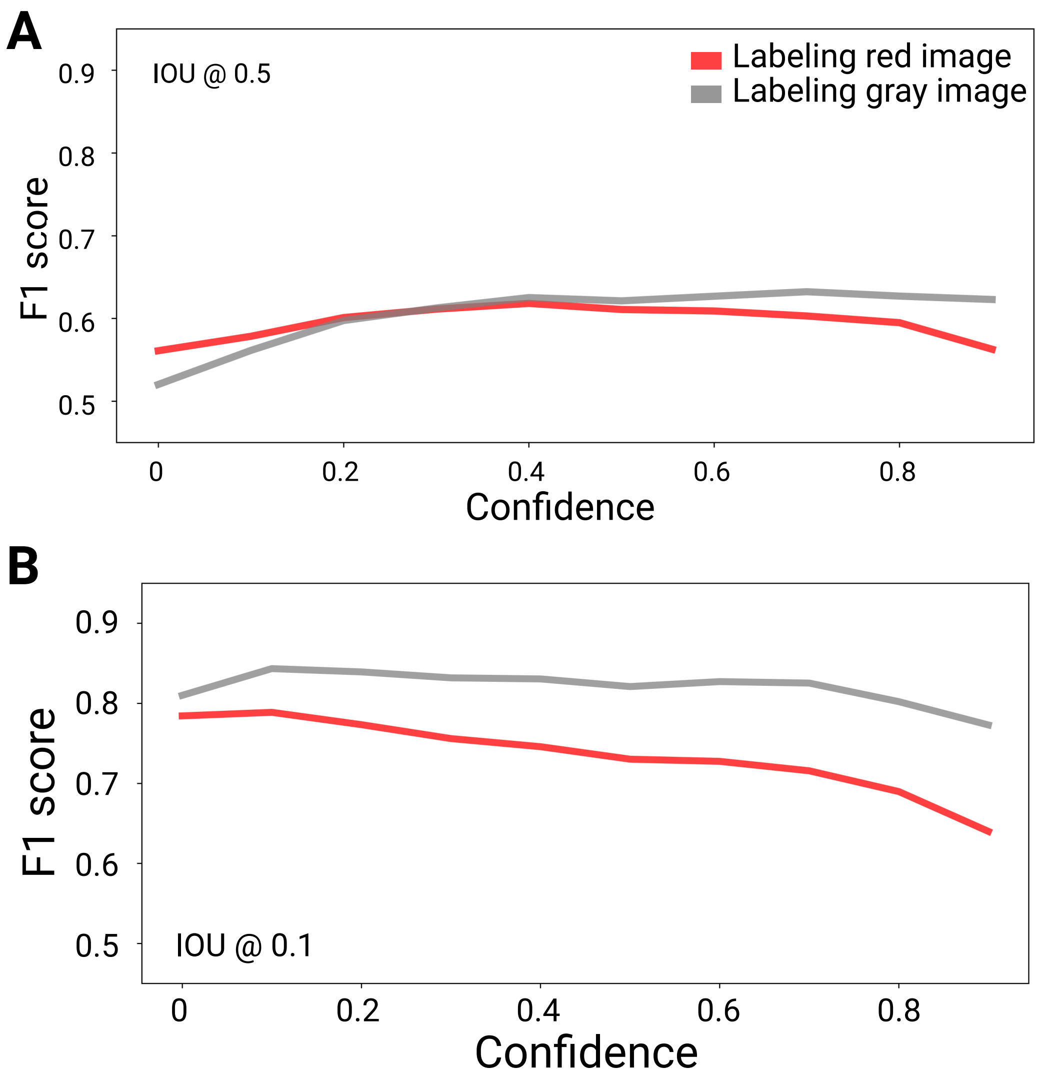


**Figure S8. H2B-mCherry images can reliably detect, and sufficiently segment MN.** A line plot depicts F1 scores upon variations to MN confidence, as per MN segmentations produced from the red mCherry images, or gray-scaled images. F1 scores were calculated, as per IOU thresholds of 0.5 for segmentation quality (A), and 0.1 object detection (B), respectively.

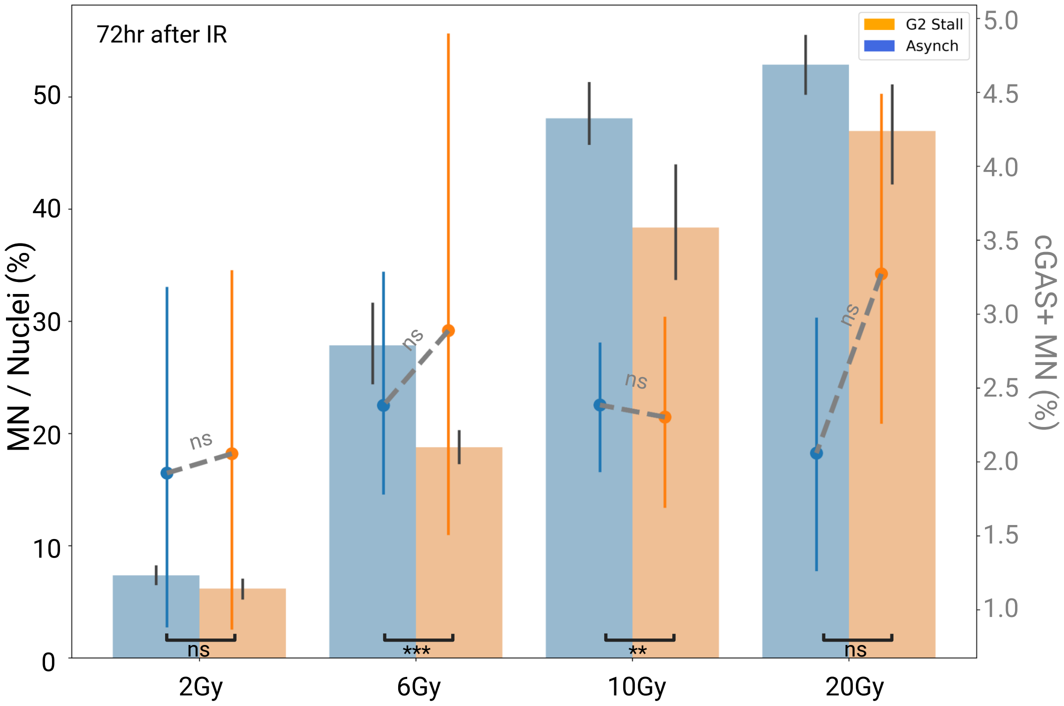


**Figure S9. MicroNucML can segment MN, useful upon its integration with cGAS-mCherry localization, 72 hours post irradiation (IR).** The barplot which aligns with the left axis and black statistical results, demonstrate the MN counts relative to nuclei counts. The point plots, which depict the proportion of cGAS+ MN, aligns with the right axis and gray statistical results (Mann-Whitney U test). Asynchronized and G2 stalled cell populations are in blue and orange, respectively. Error bars show the 95% confidence intervals.

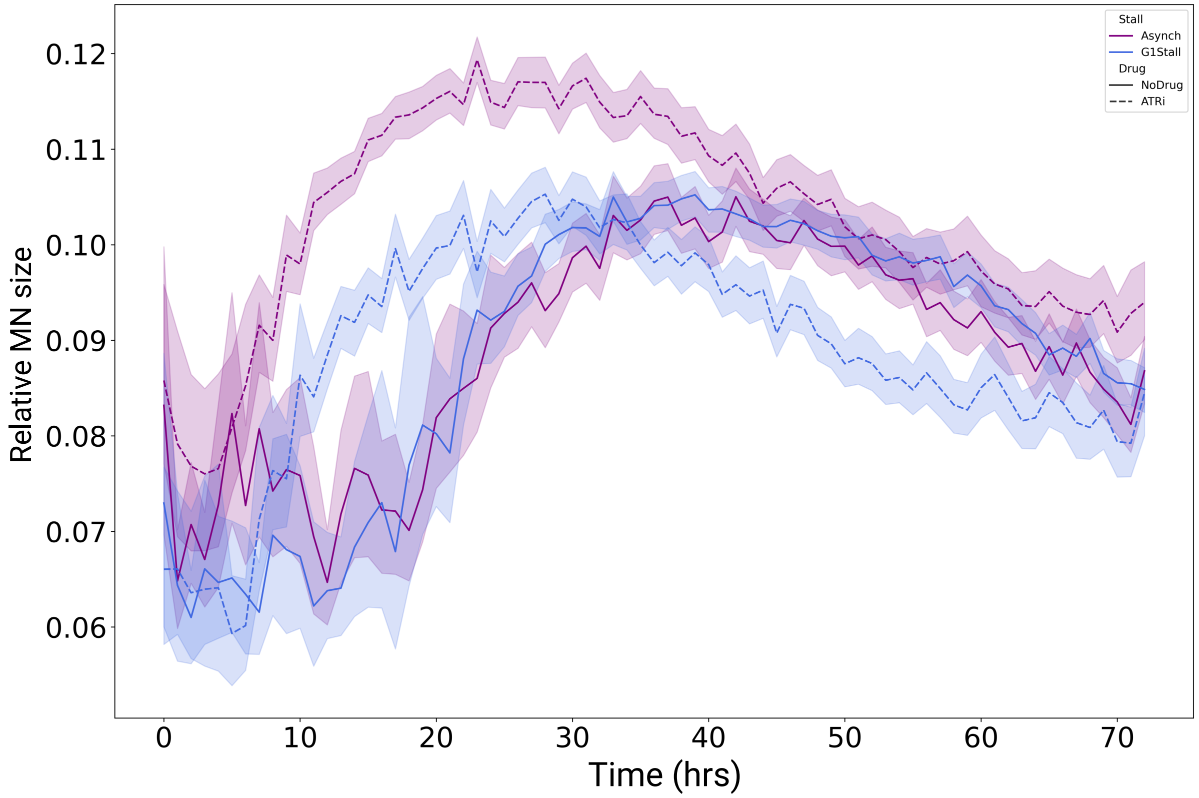


**Figure S10. The integration of MN and nuclei segmentation, at-scale, can refine biological interpretations of MN production, specifically with regards to relative MN size.** A time-course experiment, following conditions of IR, and/or ATR inhibition and/or G1 stalling. The line plot depicts the mean relative MN sizes, as per parent nuclei size. Shading describes the 95% confidence interval for a given time point. Asynchronized and G1 stalled cell populations are in purple and blue, respectively. Cells without and with ATRi exposure are show with solid and dotted lines, respectively.
